# *E. coli* transcription factors regulate promoter activity by a universal, homeostatic mechanism

**DOI:** 10.1101/2024.12.09.627516

**Authors:** Vinuselvi Parisutham, Sunil Guharajan, Melina Lian, Hannah Rogers, Shannon Joyce, Mariana Noto Guillen, Robert C. Brewster

## Abstract

**Transcription factors (TFs) may activate or repress gene expression through an interplay of different mechanisms, including RNA polymerase (RNAP) recruitment, exclusion, and initiation. TFs often have drastically different regulatory behaviors depending on promoter context and interacting cofactors. However, the detailed mechanisms by which each TF affects transcription and produce promoter-dependent regulation is unclear. Here, we discover that a simple model explains the regulatory effects of *E. coli* TFs in a range of contexts. Specifically, we measure the relationship between basal promoter activity and its regulation by diverse TFs and find that the contextual changes in TF function are determined entirely by the basal strength of the regulated promoter: TFs exert lower fold-change on stronger promoters under a precise inverse scaling. Remarkably, this scaling relationship holds for both activators and repressors, indicating a universal mechanism of gene regulation. Our data, which spans between 100-fold activation to 1000-fold repression, is consistent with a model of regulation driven by stabilization of RNAP at the promoter for every TF. Crucially, this indicates that TFs naturally act to maintain homeostatic expression levels across genetic or environmental perturbations, ensuring robust expression of regulated genes**.

---

Transcription factors (TFs) are crucial determinants of gene regulation, functioning through a myriad of regulatory mechanisms to ensure precise control of cellular processes. TFs can bind to specific DNA sequences, often located near the genes they regulate, and either promote or inhibit transcription. The regulatory mechanisms employed by TFs are diverse and complex, and regulation may alter the rate of one or more steps in the multi-step process between RNA polymerase (RNAP) binding and promoter clearance. (*1–4*). The complexity of predicting a specific TF’s regulatory function in various genetic and physiological contexts arises from the propensity of TFs to regulate multiple steps of the transcription process, combined with the intricate interplay of the number and types of TFs, their binding strengths, binding site locations, and the promoter strength (*5–8*). In particular, TFs are usually classified based on their net regulatory function rather than their mechanisms of regulation. Due to this classification scheme, it can seem surprising that same TF can have both activating and repressing interactions with different promoters, even in very similar contexts. Examples of such have been seen in both prokaryotic (*9–11*) and eukaryotic systems (*12*). Here we measure the relationship between TFs and promoters in a controlled way in *E. coli*. We systematically alter constitutive expression levels of a promoter through several methods including perturbations to basal promoter sequence, as well as by perturbing physiological conditions such as growth media or availability of polymerase. Each of these perturbations alters the constitutive expression rate of the promoter while holding other TF-related features (such as TF identity, binding site position, and sequence) constant, thereby allowing us to confidently measure the relationship between TF function and promoter identity.

## Gene regulation model to interpret TF-promoter relationship

To interpret the relationship between regulation and promoter strength, we use a simple model of gene expression that we and others have proposed previously (*13–17*). In this model (Fig. 1a), regulation by a single TF is coarse-grained into activity on two steps of the transcription process. The first mechanism of regulation affects RNAP recruitment and stability at the promoter and is parameterized by *β* (which is confined to be a positive, real number); *β* > 1 corresponds to TFs with positive stabilizing or recruiting interactions with polymerase and *β* < 1 corresponds to negative interactions with polymerase through effects such as steric hindrance. The second mechanism alters transcription initiation and promoter clearance and is parameterized by *α* (again defined as a positive, real number). Once again, *α* > 1 corresponds to TFs that increase the rate of transcription initiation while *α* < 1 corresponds to TFs that inhibit or slow the rate of this process. As such, parameter values > 1 represent activation of one step of the transcription process and values < 1 represent repression of that step. Historically, *in vitro* approaches studying mechanisms of TF regulation have tried to interpret the primary mechanism of function for specific TFs. A common model for the function of proximally-binding repressors is that of steric-hindrance where TF-binding occludes the binding of RNAP to the promoter (*β* < 1 in our model). Various *in vitro* studies have identified steric hindrance as the mechanism of action for common repressors (*18–20*) including LacI (*21–23*).

**Figure 1:**
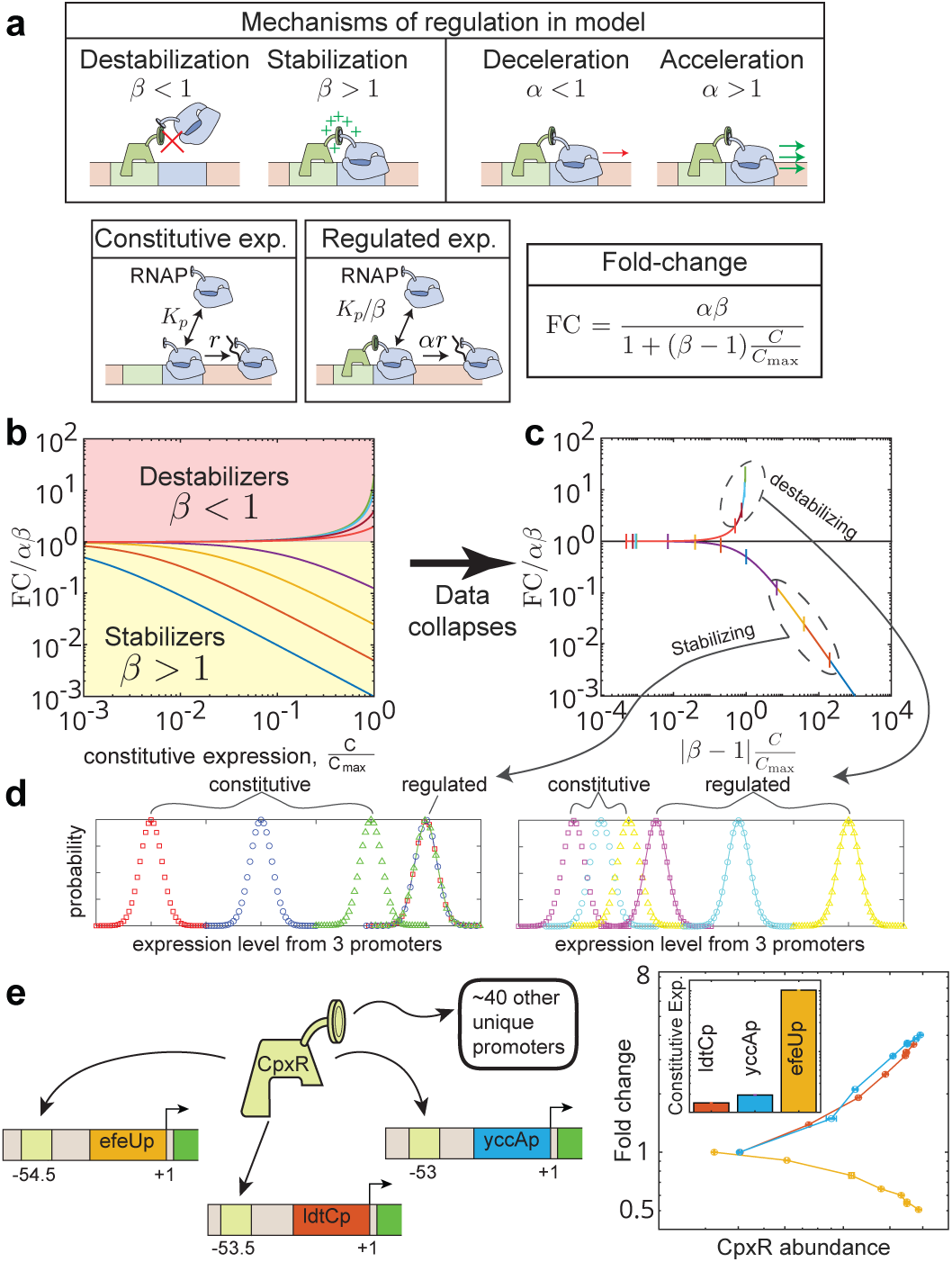
Modeling promoter dependence of TF function. (a) A simple model of TF function predicts a distinct relationship between TF function and promoter strength that depends on the nature of interactions between the TF and RNAP. (b) TFs with stabilizing interactions (*β* > 1) produce lower fold-changes on stronger promoters, while TFs with de-stabilizing interactions (*β* < 1) show the opposite. (c) The relationship collapses into two distinct curves for *β* > 1 and *β* < 1 when constitutive expression and fold-change are rescaled by the TF regulatory interactions. (d) Strong stabilizing interactions will buffer changes to constitutive expression while destabilizing interactions amplify them. (e) The TF, CpxR acting on three natural promoters demonstrates that regulatory function can have a strong dependence on the strength of the regulated promoter.

We make no assumption, *a priori*, about how any TF will function; instead our model enables interpretation of TF function *in vivo* through these two mechanisms (*12, 13, 15, 24*). In the limit of saturating TF concentrations, the model makes a simple prediction for the relationship between fold-change in gene expression and constitutive promoter strength, *C* (Fig. 1a) where fold-change is the ratio of expression at saturating TF levels (regulated, R) to the expression in the absence of that TF (constitutive, C). This relationship is plotted for several values of *α* and *β* in Fig. 1b. Essentially, if a TF is stabilizing (TF-RNAP interactions are favorable, *β* > 1), a TF will enact lower fold-change on stronger promoters. That is to say that repressors will repress more and activators will activate less on strong promoters compared to weaker ones. The opposite relationship is predicted if the interactions are destabilizing (TF-RNAP interactions are unfavorable, *β* < 1). Crucially, because there are two possible mechanisms of regulation in our model, a destabilizing TF can still activate and a stabilizing TF can repress, contingent on the TFs role on the other step of transcription.

Crucially, Fig. 1c demonstrates a stringent prediction from our model for TFs that regulate through positive TF-RNAP interactions (*β* > 1): when the basal promoter is strong, fold-change will scale with the inverse of constitutive expression level (*C*^−1^). Importantly, this corresponds to a behavior where stabilizing TFs will elicit the same regulated levels of expression independent of the constitutive expression of the promoter (Fig. 1d). Alternatively, destabilizing TFs will drive divergent levels of regulated expression from promoters with relatively small differences in constitutive levels.

As an example of the complex dependence between TF regulatory function and promoter identity, consider the TF CpxR which regulates more than 40 different promoters in *E. coli* (*25*). Fig. 1e shows data for the fold-change as a function of CpxR concentration for three CpxR-regulated promoters (ldtCp, yccAp, and efeUp). These promoters have CpxR binding sites at approximately the same position relative to their transcription start site (TSS). However, two of these promoters are activated by CpxR (ldtCp and yccAp) while the other (efeUp) is repressed. Although some context-specific details (such as TSS, 5^′^ untranslated region, ribosome binding site, etc.) are different between these three promoters, one notable difference is that each core promoter gives rise to significantly different constitutive (unregulated) expression level. The inset to Fig. 1e shows the measured expression of each promoter in a CpxR knockout strain. CpxR activates the two weak promoters and represses the stronger promoter, which has 100−fold higher constitutive expression. Previous studies have suggested that CpxR acts primarily through positive, stabilizing interactions with RNAP (*β* > 1) (*10*), which is qualitatively consistent with the trend seen in our data in Fig. 1e and with our prediction that a TF can switch from activating to repressing gene expression dependent on the core promoter strength. We do not require that the intrinsic function of the TF changes or is “context specific”, it is a basic expectation of our model. However, there are many uncontrolled aspects of this measurement; the TF binds to slightly different positions on the promoter, the binding site sequences are different, and there may be differences in other regulatory factors involved at each endogenous promoter. Hence, we sought to measure the dependence of regulatory function on promoter strength in a system that controls for these contextual confounds.

## Measuring the relationship between TF function and promoter identity on a synthetic promoter library

We chose eight different TFs identified from a previous study (*15*) to examine the relationship between their regulatory effect and the activity of the regulated promoter. These TFs were chosen due to clear evidence of regulatory interactions on a synthetic promoter cassette which contained only a single binding site for the TF (*15*) (details in Table S1 and Fig. S1c). An overview of the experimental method is outlined in Fig. 2a. To create a library of promoters with a spectrum of constitutive strengths, we mutated the −35 region of the promoter (153 possible combinations of single and double mutations in the −35 regions Fig. S1a,b) and randomly sampled 96 different clones. This library typically showed a range of constitutive expression levels varying from 100 to 1000 fold relative to the weakest promoter. We measured the expression of each promoter variant both in our library of inducible TF strain (*24*) at full induction (*T F*^++^), and in a strain where the TF has been deleted (*T F*^−^). We then plot the measured fold-change of each promoter against the constitutive expression level of that promoter (Fig. 2b).

**Figure 2:**
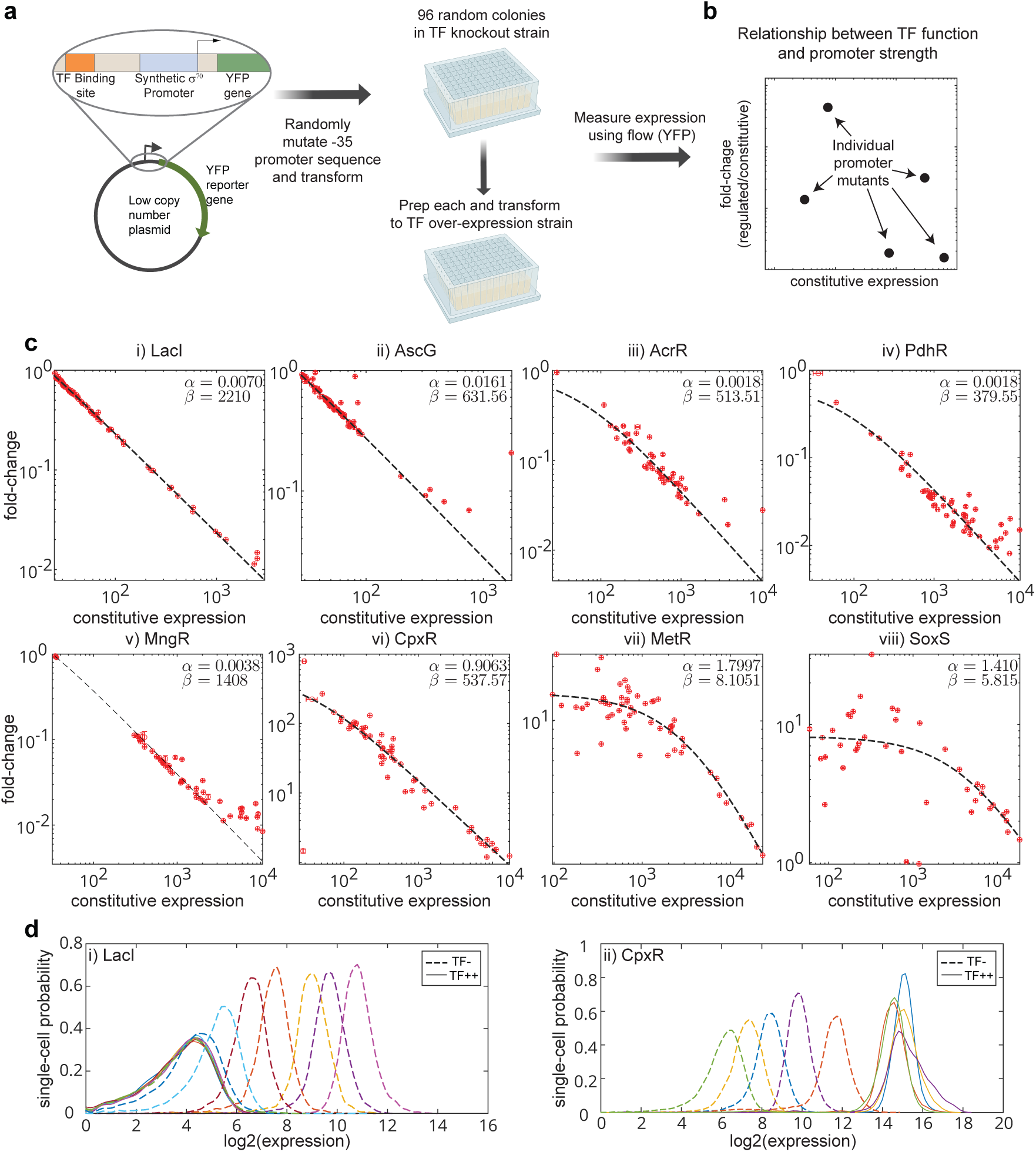
Measuring the relationship between TF function and promoter activity for synthetic promoter variants. (a) Experimental approach to measure fold-change and constitutive expression levels of random promoter variants. (b) Each data point in our plots represents the constitutive and regulated level of expression for one promoter variant. (c) For 8 TFs measured here, the relationship between TF function and promoter strength conforms well to the predicted scaling of strong stabilizing interactions (black dashed lines). Subplots (*i*)-(*iv*) show TFs binding downstream of the promoter, while (*v*)-(*viii*) show TFs binding upstream of the promoter. Furthermore, panels (*i*)-(*v*) show TFs considered to be repressors and (*vi*)-(*viii*) show activators. (d) As expected from the theory for stabilizing TFs, the regulation by both CpxR and LacI cause a robust level of regulated expression from promoters with diverse unregulated levels of expression.

Fig. 2c shows the data for the eight TFs: five repressors (LacI, MngR, PdhR, AscG, and AcrR) and three activators (CpxR, MetR, and SoxS). The top row of regulatory interactions is for TFs with binding sites downstream of the promoter (centered at positions ranging between +11 to +16.5 from the TSS) and the bottom row features regulatory interactions from TFs with binding sites upstream of the promoter (centered at positions between −52.5 to −64 from the TSS); see Table S1 for details. The dashed line in each plot is a fit to the theory in Fig. 1a with *αβ* and *β* as fit parameters. The maximum promoter activity, *C*_max_ is set equal to the level of the promoter with the maximum expression in each dataset. By setting *C*_max_ this way, we establish a lower bound on the maximum possible expression. Intuitively, if *C*_max_ is increased, the value of *β* increases proportionately (see Fig. S1e,f). Crucially, each of these eight TFs shows the predicted scaling relationship for stabilizing TFs independent of function (activation or repression) or binding location on the promoter. However, for two activators, MetR and SoxS, the data for the weakest promoters fall into the regime where FC is constant with promoter strength consistent with weaker stabilizing interactions for those TFs as measured by lower values of *β* from the fit.

A widely accepted model of repression used by us and others in the past, is regulation purely through negative interactions with RNAP (steric hindrance) (*6, 10, 26, 27*). In our general model, this corresponds to setting *β* < 1 and *α* = 1. In this case, the model specifies that the fold-change at low promoter strength will equal to *β* and the fold-change of strong promoters will increase and eventually will be equal to 1 for the strongest promoters; our data shows that neither of these holds true. This goes against the long-held steric hindrance model which fails to describe every repressive dataset in our study (Fig. 2c (i)-(v)). In our model, *β* > 1 is a requirement to capture the observed inverse scaling of fold-change with promoter strength. Therefore we conclude that there is a common positive interaction between TF binding and RNAP availability at the promoter in every regulatory interaction measured here and that the net regulatory effect is determined by the magnitude of impact on the second step (*α* in our model). Importantly, this positive, stabilizing interaction may arise through direct interactions with RNAP or indirect mechanisms such as changing the DNA’s local propensity to bind RNAP.

An important consequence of the stabilizing relationship seen for most TFs tested is that as constitutive expression levels change, the regulated levels remain constant for TFs with relatively high values for *β*. This is demonstrated in Fig. 2d for two TFs, LacI and CpxR. We plot the single-cell distribution of constitutive (dashed lines) and regulated (solid lines) expression for several promoters sampled from the a range of constitutive expression levels in the library where each color represents a unique promoter sequence. The constitutive expression levels vary roughly 200-fold over these promoters. However, the corresponding measurement of the regulated expression levels (solid lines) vary by only roughly 2-fold. Thus, due to this inverse scaling relationship between TF function and promoter activity, the TF acts to buffer changes to expression from induced mutations in the promoter for both an activator and a repressor.

## Measuring the relationship between TF regulation and promoter strength in different physiological conditions

The effect of physiological perturbations on constitutive expression has been studied using simple models that demonstrate the scaling between growth rate and constitutive expression (*28, 29*). Despite strong coupling between growth rates and transcription regulation little is known about how different TF functions are influenced by physiological perturbations (*29*). So, we measured the relationship between TF function and promoter strength by inducing changes to constitutive expression levels through physiological perturbation of growth rates using an array of carbon sources. We measured these effects using two TFs: LacI and CpxR. We measured each promoter mutant library in six different minimal media conditions supplemented with a range of carbon sources that yielded doubling times between 55 (glucose) and 230 minutes (acetate). We expect significant changes to several global regulators including the RNAP concentration between these media conditions that impact the basal strength of all the promoters. However, the TF concentration is also perturbed by these growth conditions; most notably the TF concentration in the cell increases significantly in the slowest conditions (pyruvate and acetate). To ensure the validity of our hypothesis testing, it is essential that fold-change is either measured at a saturating TF concentration, where further increases in TF concentration do not affect the fold-change or that the TF concentration is kept consistent across all conditions (Fig. S1d). We find that the changes in TF levels do not perturb the quantitative level of LacI regulation because it is already saturating for our binding site affinity (*i.e.* increasing TF concentration does not alter fold-change, cyan data points in Fig. S1c), whereas CpxR is at sub-saturating concentrations and the regulation changes with TF concentration (brown points in Fig. S1c). Thus, for CpxR, we do not include data from pyruvate and acetate in which the TF concentration is significantly different. Including those data points (Fig. S2b) shows the same scaling relationship but with a larger *y*-intercept corresponding to increased activating function across all promoters, consistent with the theory in Fig. S1d.

Perturbing the growth rate changes the fold-change from the promoter library, with slower-growth rate systematically showing lower fold-changes, however, the scaling relationship is preserved across the entire dataset for both TFs (Fig. 3a). Furthermore, the relationship between the regulation of any one promoter across all growth rates shows this same scaling (see inset to Fig. 3b); Fig. 3b shows the distribution of slopes from a straight-line fit to each promoter across the measured conditions for both TFs. Most promoters have a scaling relationship with a slope of −1. Importantly, this scaling is seen across individual promoters with different growth rates as well as across the collection of promoters at a single growth rate and this relationship is invariant between genetic or physiological changes for both activator and repressor. In Fig. 3d we show the single-cell expression distribution for a single promoter across various growth rates. Once again, although the constitutive expression level of the promoter depends strongly on growth rate, regulation is largely capable of buffering these changes where the stable level of expression is set by the identity of the TF regulating it; CpxR collapses the expression from the promoter into a higher expression state while regulation by LacI also collapses the data but in a repressed state.

**Figure 3:**
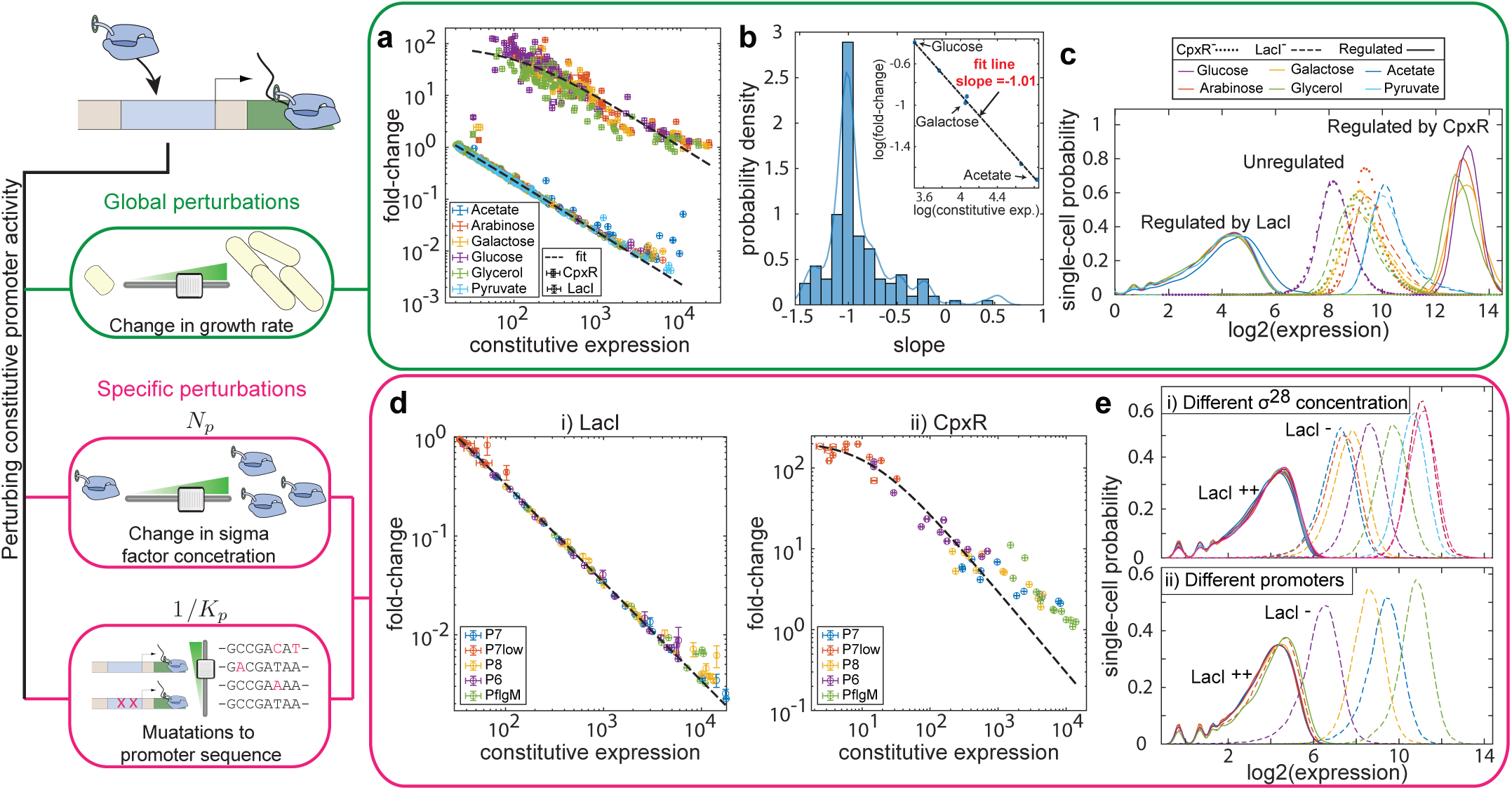
Scaling of regulation is conserved across different methods of perturbations to the constitutive expression. (a) The constitutive expression level is altered via changes in the growth rate achieved by supplementing media with different carbon sources. The plot shows the measure of regulation when the promoter mutants of LacI (circles) and CpxR (squares) were grown in media containing a variety of carbon sources (different colors). (b) Distribution of slope obtained from a linear fit of fold-change as a function of constitutive expression for any single promoter in different media. Inset shows the straight-line fit to the data for one of the promoters grown in 6 different carbon sources. (c) Representative single-cell distribution of the fluorescence of unregulated (dotted lines) and regulated expression (dashed lines) across different growth media. The global perturbations alters the growth rate and the constitutive expression levels of the library of promoters but not the regulated expression levels both for activation (CpxR) and repression (LacI). (d) The constitutive expression is altered via specific perturbations to RNAP concentration (via changes to σ^28^ concentration) or promoter sequence. Measure of regulation by LacI (i) and CpxR (ii) for specific perturbations. Each color on the plot is a different promoter and each data point is a fixed concentration of σ^28^. (e) Single-cell distribution of regulated and un-regulated expression for changing σ^28^ concentration (i), or promoter strength (ii). There is no change in the regulated expression (solid lines in each plot) for any specific perturbation to constitutive expression (dashed lines in each plot).

Once again, by global perturbation to the constitutive expression through physiological changes, we find the same fundamental mechanisms of regulation that “restore” the regulated state expression level.

## Promoter-TF relationship is predictive for regulation of alternative sigma factor promoters

We next designed a system where we could systematically perturb only the constitutive expression levels. Ideally, we want to change expression levels of our promoter by altering the availability of RNAP, however, limiting any subunit of the core σ^70^-RNAP holoenzyme induces extreme physiological changes (*28, 30, 31*), making it hard to isolate the regulation of just our one target gene. As such, we designed a system to control the availability of an alternative sigma factor protein, σ^28^ by expressing it under an inducible system. The advantage of this system is that the σ^28^-RNAP holoenzyme recognizes an orthogonal promoter sequence to the housekeeping σ^70^, which is the primary sigma factor in *E. coli* (*32, 33*). The promoters recognized by σ^28^ are involved in flagellar synthesis (*34–36*) and the mechanisms of control (anti-sigma factors) are known (*37, 38*). In this system, we have direct orthogonal control over both the physiological (via σ^28^-RNAP concentration) and the genetic component (via promoter sequence) of the constitutive promoter activity.

We designed a system where the expression of σ^28^ from the FliA gene can be induced with vanillic acid, and the endogenous copy of the gene is knocked out. We then swapped our σ^70^ specific promoter used previously with a promoter specific for σ^28^. We see that increasing the concentration of vanillic acid results in higher expression from σ^28^ promoters due to the elevated levels of active σ^28^-RNAP holoenzyme. The dashed lines in Fig. 3e(i) show the achievable range in our inducible system for one promoter with different vanillic acid concentrations. To pair with this control of σ^28^, we designed five promoters that span our measurable range of expression levels. The dashed lines in Fig. 3e(ii) show the distribution of expression for single cells from each promoter at intermediate vanillic acid concentrations. As a final means of expanding our range of expression, we make these measurements in a strain with the endogenous anti-σ^28^ factor FlgM (which competitively binds to free σ^28^ and thus lowers available σ^28^-RNAP holoenzyme availability (*39*)) as well as in a strain with FlgM knocked out.

Fig. 3d shows these measurements for regulation by a repressor, LacI(i), and an activator, CpxR(ii). Each promoter (represented by different colored points) is measured at eight different σ^28^ induction levels both with and without the endogenous expression of the anti-sigma factor, FlgM. The result is a constitutive expression range over more than 3 orders of magnitude. Importantly, the fold-change scales over this range as expected for a single set of *α* and *β* parameters for each regardless of whether genetic or physiological perturbations changed the constitutive expression level. In Fig. 3e(i-ii), we show once again how this data collapses when regulated by LacI for both physiological (top panel) and genetic (bottom panel) perturbations. Different CpxR promoters do not align perfectly with theoretical predictions; however, within each promoter type, the inverse scaling relationship is preserved. The absence of a complete data collapse for CpxR may occur due to sub-saturating TF concentration (Fig. S1d right panel) or limitation to the σ^28^ availability.

## Measuring the relationship between TF function and promoter identity in natural promoters

To this point, we have observed a specific scaling relationship between TF function and constitutive promoter strength that is pervasive in simple synthetic promoters designed to be regulated only by a specific TF. Here we extend this concept to determine the applicability of this relationship in naturally occurring promoters with complex regulatory architectures such as regulation by DNA looping, multiple binding sites for the same TF, or interfering binding sites by other TFs.

In Fig. 4a, we show measurements of several endogenous promoters regulated by the TF, SoxS (MarA and Rob, the other isoforms of SoxS are deleted from the strain to avoid any cross-talk): poxBp (yellow), decRp (blue), and fldAp (red) (*40*). These promoters were chosen because their endogenous binding sites are located at a similar position relative to the promoter (Fig. S3a). We created 96 random mutants of each promoter using the same strategy as described above for the synthetic promoters. Unlike the synthetic promoters that have minimal changes between different TFs, the natural promoters differ in every aspect including the 5^′^ UTR region, ribosomal binding site (RBS) affinity, and transcriptional start site. These differences in translation efficiency and mRNA stability make comparing constitutive expression levels across different promoters challenging. To normalize for these differences, we measured the relative mRNA and protein expression levels between each of the 3 promoters and corrected the constitutive expression by normalizing *C* by the ratio of mRNA to protein expression levels of each promoter (see Fig. S3b). Correcting for these features helps reduce the non-transcriptional differences across the three promoters to compare their transcriptional activity directly. Crucially, the effect of this multiplicative correction factor amounts to translation of the data horizontally along the *x*-axis and does not influence the scaling of any one dataset.

**Figure 4:**
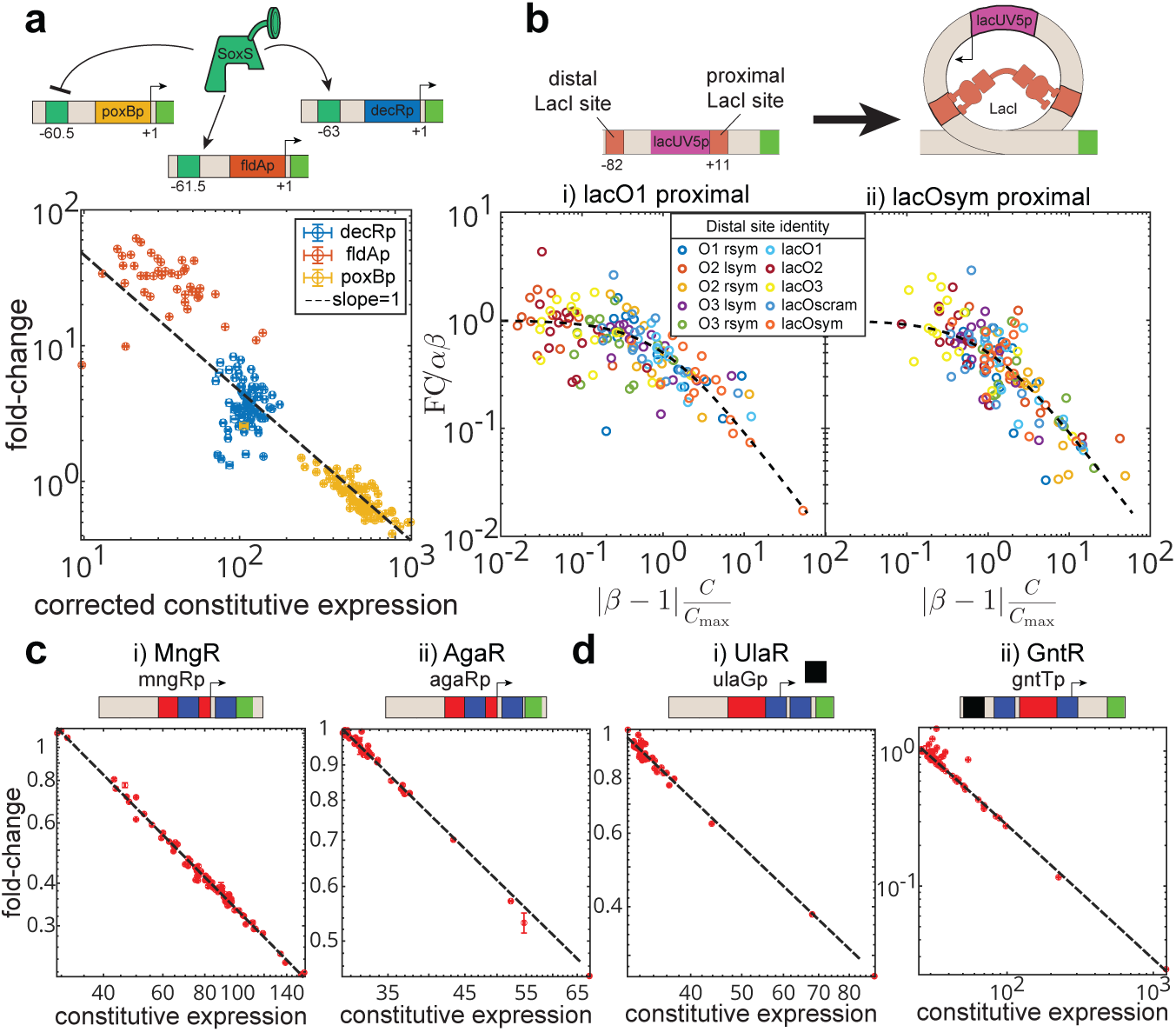
Promoters with complex regulatory architecture show stabilizing relationship with mutant promoter library. (a) The top panel shows three native promoters of *E. coli* regulated by SoxS. The relative position of the binding site is shown as a complete green square with the center of the binding site marked underneath. The bottom panel shows the plot of fold-change against the corrected constitutive expression level for the three native promoter mutant libraries. (b) The top panel is a schematic representation of the DNA looping mediated regulation by LacI. The bottom panel is the massively parallel reporter assay data from (*43*) plotted using un-induced/induced as a proxy for the fold-change and induced expression as the constitutive expression. Data corresponding to the strong proximal binding sites (LacO1(i) and LacOsym(ii)) and all of the 10 distal binding sites are plotted. (c) Measure of regulation for a promoter architecture with two binding sites for the TF of interest (dark blue squares) around the promoter (red square). (D) Measure of regulation for promoters with binding sites for other TFs (black squares) in addition to the binding site for the TF of interest.

The collection of promoter mutants shows regulation of up to 70-fold activation down to 3-fold repression between the three promoters. Each promoter shows a good agreement with the predicted scaling relationship of stabilizing interactions. The relationship is nearly universal between the three promoters across the entire range of fold-change, however, the fold-change values for the fldAp promoter are systematically 4−fold higher. This may be due to slightly stronger regulatory interactions from SoxS at −61.5 compared to the other two positions (*15*) or perhaps due to slight differences in binding site affinity of SoxS across the three unique binding sites. While measuring binding affinity is beyond the scope of this paper, we assessed the proximity of the annotated binding sites on decRp, poxBp, and fldAp to the SoxS consensus binding site using the TF PSSM browser in RegulonDB (*40*). This tool conducts a MEME analysis and generates p-values for the annotated binding sequences. Lower p-values signify sequences that closely resemble the consensus, whereas higher p-values indicate a greater similarity to a random sequence. We expect the SoxS binding site of decRp and poxBp to be of similar strength (p-values of 1.14 × 10^−3^ and 2.74 × 10^−3^ respectively), while the site in fldAp is significantly closer to the consensus (p-value 1.76 × 10^−4^) (*41*). Converting distance from consensus sequence to affinity is not straightforward, but this qualitatively supports the cause of the shift in fldAp data. Furthermore, we highlight that SoxS can switch roles from activation to repression just by changing the strength of the core promoter. It is noteworthy that SoxS has been previously thought to be involved in stabilizing RNAP by acting as a co-sigma factor (*42*), consistent with our data. However, the ability to repress strong promoters implies that it must also repress the initiation or a further downstream step of transcription.

We next explored the common natural regulatory architecture of DNA looping, where a TF binds to two sites on the promoter simultaneously to repress expression. Yu *et al.* (*43*) examined regulation by LacI with any one of ten different LacI binding sequences at both the distal and proximal binding location using a collection of promoter sequences. In Fig. 4b we show the data with the two strongest sites (O1 and Osym) at the proximal position and any of the 10 different sequences at the distal position. The spacing between these two LacI binding sites is set to mimic the natural distance between O1 and O3 in the *lac* operon. Since distal site choice will influence the level of regulation and thus the parameters of the model, we fit each data set to the theory and plot all of the data according to the rescaled axes suggested in Fig. 1c. As can be seen, the data from every examined looping architecture collapses to a single curve and obeys the expected scaling relationship observed for stabilizing interactions. This same relationship holds even for the weaker proximal sites, although the regulation gets very weak (Fig. S5). Interestingly, one trend that can be seen in this data as predicted by our theory, the stronger the proximal site, the more the data skews to the “strong regulation” part of the universal plot which scales with the inverse of the constitutive expression(Fig. S5).

We further examined other TFs acting on endogenous promoters with more complex regulatory features where we have no insight into the mechanisms of regulation. Fig. 4c shows two situations where regulation of the endogenous promoter occurs through two binding sites: (i) mngRp regulated by MngR and (ii) agaRp regulated by AgaR. Although naturally these promoters are wired for feedback, since we have deleted the endogenous TF gene, our measurements isolate the regulatory role for fixed, saturating TF concentration. Furthermore, Fig. 4d shows two other natural promoters regulated by TFs we have previously studied (UlaR, GntR (*15*)). These promoters also have two binding sites for the measured TF interaction as well as additional TF binding sites; ulaGp features IHF regulation at an unknown location and gntTp features a CRP binding site. Again, in all cases, we find the same scaling relationship between fold-change and constitutive promoter strength. Interestingly, these promoters which are regulated by other TFs, show decreased response to promoter perturbations compared to synthetic promoters designed to be regulated by only a single TF. This is in line with the idea that TFs are capable of stabilizing such genetic perturbations.

## All data collapses to the same scaling law

To demonstrate the universal nature of this TF-promoter relationship in our measurements, in Fig. 5 we collapse all the data from this study onto the same plot by rescaling the constitutive expression of each data set by (*β* − 1)*C*/*C*_max_ and the fold-change by 1/(*αβ*) akin to our theory plot in Fig. 1c. The data shows a strong, universal collapse to the stabilizing TF theory predictions. Furthermore, we add previously published data sets: from the activator AraC (*44*), from LacI looping (*43*), from the activator CRP (*45*), and from *in vitro* measurements of the TF, CarD from the bacterium *Mycobacterium tuberculosis* (*46*). Strikingly, across nearly 6 orders of magnitude in the fold-change split between activation and repression (see inset to Fig. 5), the relationship between TF function and constitutive promoter strength collapses to a single functional form. This includes data from genetic perturbations, physiological perturbations, and *in vitro* measurements by a different group. In all cases, the data conforms to a picture of TF regulatory function that has a conserved regulatory mechanism.

**Figure 5:**
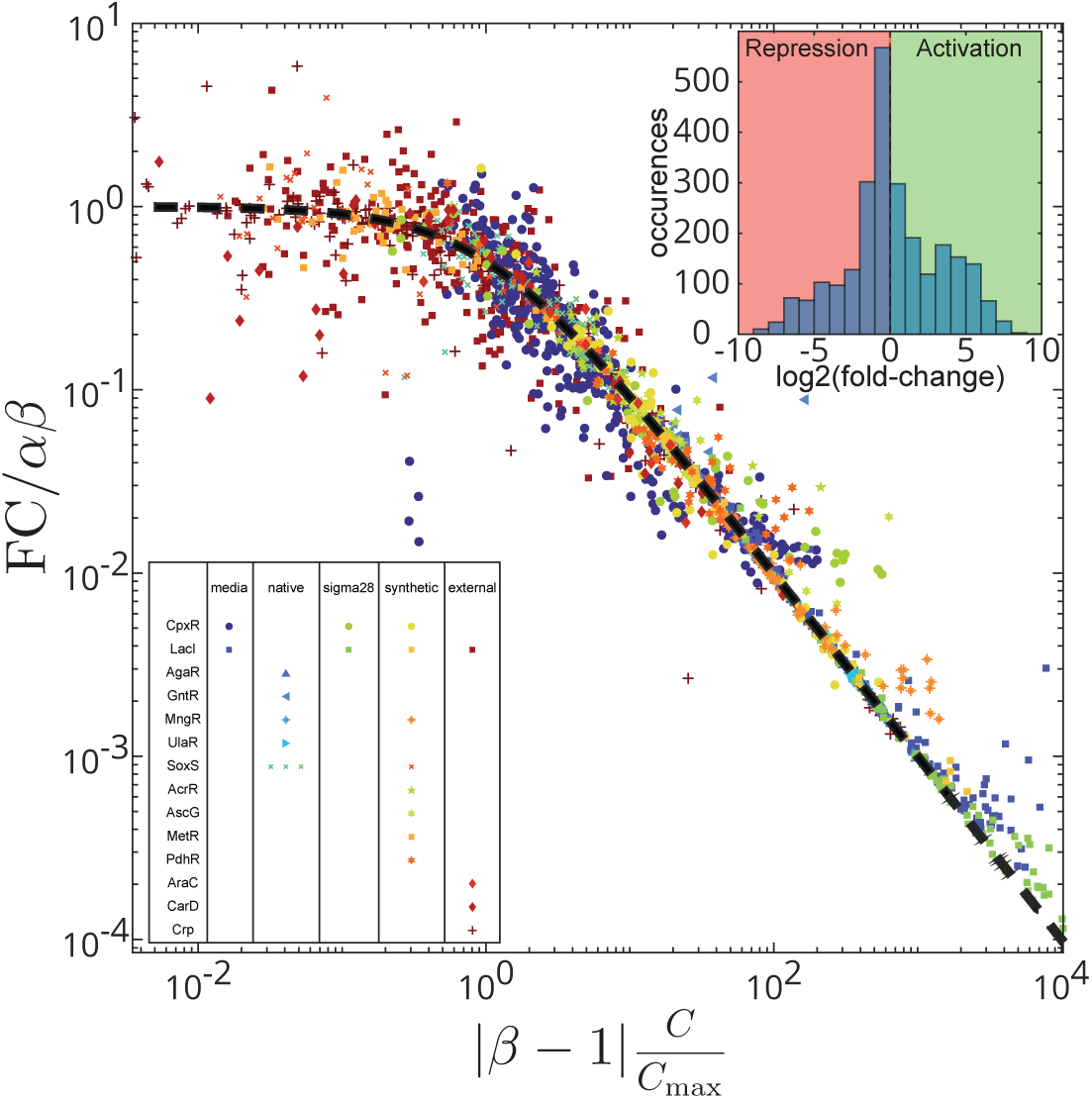
Universal collapse for all regulation data. All data from Figs. 2− 4 are renormalized using the *α* and *β* obtained from each fit. We also include data from refs. (*43, 44, 46*) which we fit the same way as described above for *α* and *β* values. The black line represents the zero-parameter theory line: *f* (*x*) = 1/(1 + *x*). All data collapses to a single theory curve suggesting a conserved universal mechanism of action between all measured regulation. The inset shows a histogram of fold-change values for all points included in the figure.

## Discussion

The relationship between TFs and the promoters they regulate is an important determinant of gene expression levels. Any given TF may regulate a battery of promoters with distinct features and these promoters will be regulated in various physiological conditions. Therefore, an understanding of the principles that determine how these features govern regulated gene expression is crucial for determining how gene expression changes upon mutation or environmental conditions and how these affect physiology. Using *in vivo* measurements in *E. coli* combined with a model of gene regulation, we discover a universal principle that TFs function to stabilize RNAP at promoters and the fold-change in expression enacted by strongly-acting TF scales as the reciprocal of the constitutive activity of the promoter. This function applies to all TFs tested, both activators and repressors, acting on both synthetic and naturally occurring promoters. The relationship holds if we perturb the “promoter strength” through (1) random genetic mutations to the promoter (2) physiological perturbations to the growth rate that change the propensity of transcription, (3) through direct perturbation of RNAP availability, or (4) any combination of these perturbations.

It is particularly surprising that every repressor measured here shows this same “stabilizing” relationship with promoter strength. This is contrary to what is often considered the common model of repressor action: steric occlusion prevents RNAP from binding to the promoter in the presence of a repressor. This is sometimes termed the “competitive model” of TF function. This idea has been supported in some *in vitro* studies specifically for the function of LacI (*22, 47*). This model is intuitive since LacI binds very close to the *lac* promoter, with the 5^′^ end of the binding site at +1 on the promoter. However, there are conflicting *in vitro* studies which have suggested regulation by LacI occurs at the step of transcription initiation (*48–50*). Straney and Crothers specifically found that LacI increased RNAP binding by more than 100−fold in their *in vitro* assays (*48*), although these contradictory results have sometimes been dismissed as an artifact of low ionic concentrations in the *in vitro* experiments (*22, 51*). Our *in vivo* findings support both findings by Straney and Crothers: (1) LacI negatively regulates at the step of initiation and (2) LacI positively affects stability of RNAP at the promoter. In fact, we find this to be a general regulatory feature of TFs across every interaction measured in our study. Surprisingly, our study does not find any evidence for regulation by steric occlusion, as inferred from our model.

An important feature of stabilization interactions is that regulation is “restorative” in nature. Perturbations to the base level of gene expression, either through mutations to the promoter region or changes to the physiological state which may up or down-regulate global expression rates, will be compensated when the TF-promoter has a stabilizing relationship. On the other hand, destabilizing TFs, which we do not find to be common, would exacerbate perturbations to base expression levels. A speculative explanation for the prevalence of stabilization relationships observed could be that the robust relationship between TF and promoters is evolutionarily favored; A TF whose function compensates for perturbations will help make expression levels robust and help maintain homeostasis of the cell.

This observation of a conserved mode of regulation highlights the vapidity of the labels “activator” or “repressor” as a characterization of TF function and highlights the importance of understanding the mechanisms of action that affect regulation. If it is the case that most TFs operate through stabilization, then all repressors are inherently “incoherent regulators” (*10, 12–15, 52*) and their identity as a repressor depends entirely on the basal strength of the regulated promoter.

## Acknowledgments

We wish to thank Marian Walhout, Manuel Razo-Mejia, and Griffin Chure for helpful discussions.

## Author contributions

## Data and materials availability

All data is available in supplementary files.

## Supplementary Materials

### Materials and Methods

#### Mathematical relationship between fold-change and constitutive expression

Thermodynamic models of gene regulation are widely used to obtain quantitative gene expression predictions using a minimum number of unknown variables. One common approach to reducing the number of unknown variables in thermodynamic models is to assume that the promoters are weak (*6, 26, 27*). We recently revisited the thermodynamic framework for its general applicability by relaxing the weak promoter assumptions (*14, 15*). TFs interact in different ways including directly binding to the DNA or RNAP or to a co-factor, to bring forth a regulation (*11*). Rather than attributing a specific regulatory function to the TF, it is modeled based on two key phenomenological parameters: recruitment (quantified as *β*) and acceleration of RNA polymerase (quantified as *α*). This assumption serves to consolidate the regulatory role of TFs, which is often debated as context-dependent activators or repressors. The thermodynamic framework for gene regulation thus reveals an elegant relationship between promoter strength and the maximum expected regulation by a TF (fold-change, FC), as outlined in (*14, 15*). A simplified version of the relationship for constitutive expression, C, and regulated expression at saturating TF, R is represented in Equation (1-3) (Refer to SI for finite TF concentration).

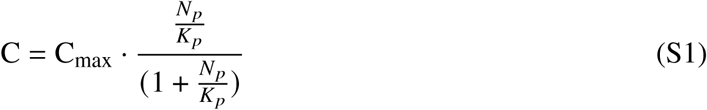

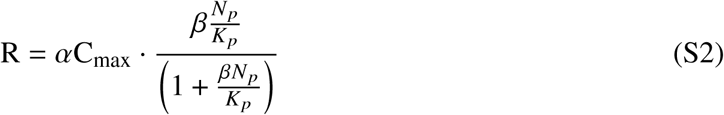

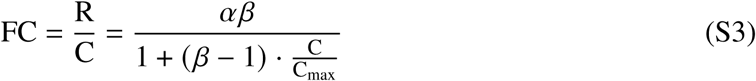

where FC is the fold-change measured as the ratio of expression in the presence of TF to the expression in the absence of TF which is also the constitutive expression, C. Thus, both FC and C, are readily measurable quantities.

#### Bacterial Strains

All strains used in this study are based on the collection of TF library strains as in (*24*) and synthetic reporters are from (*15*). Reporters with native promoters are constructed by cloning the promoter sequence (starting from the transcription start site and moving upstream with all the regulatory regions included) upstream of the YFP reporter. For sigma factor titration, the regulator *vanR* and the promoter, vanCp are PCR amplified from the marionette strains (*53*) and combined by splice overlap extension PCR to a spectinomycin cassette and inserted by lambda red recombinase into the native locus of *fliA* (the gene for σ^28^ sigma factor) replacing the native fliAp promoter with vanCp. The regulator *vanR*, is inserted bidirectionally to *fliA* gene. The *λ* prophages carrying Spec^R^::vanCp::fliA is transduced into the control and library strains of CpxR and LacI and selected for spectinomycin resistance. To introduce the anti-sigma factor knockout, the *λ* prophage carrying the *flgM* knockout from the Keio collection is first transduced into the corresponding control strains, selected for kanamycin resistance, and then the kanamycin cassette is cured by the use of pCP20 plasmid. The *λ* prophages carrying the TF construct (library strain) and the Spec^R^::vanCp::fliA constructs are then sequentially transduced into the control strains with the *flgM* knockout, via two tandem P1 transduction.

#### Creation of mutant library

Oligo pools homologous to the −35 region of the promoter are designed by replacing two of the wildtype sequences with random nucleotide “N” as depicted in Fig. S1b). 15 oligos are designed for a −35 sequence resulting in 154 unique combinations of the −35 sequences. For DL5p, the −35 region is selected from the literature. For native promoters, the −35 sequence is either based on the predictions in regulonDB (*40*) or based on the Salis promoter calculator (*54*). For synthetic DL5p constructs with the binding site located downstream of the promoter or for the native promoters, the reporter plasmids are PCR amplified with primers that target regions immediately upstream and downstream of the −35 region of the promoter and the single-stranded oligo pool with the mutations is used as a bridge to complete the plasmid Fig. S1a. The amplified PCR products are digested with DpnI enzyme, purified, assembled using NEB Hifi DNA assembly reaction, and transformed to corresponding control strains where the TF of interest is knocked out. For synthetic DL5p with binding sites located upstream, the forward primer amplifying the plasmid carries mutations for the −35 region at the 5^′^ end and the reverse primer is immediately upstream of the −35 region. The plasmid is then blunt-end ligated using NEB’s KDL reaction mix and transformed to corresponding control strains where the TF of interest is knocked out. 96 different clones per mutant library are sequenced to exclude any mis-assembled constructs and the rest of the constructs are transformed to the corresponding library strain for further measurement.

#### Transformation to the library strains

96 independent colonies per mutant library are inoculated in 1 mL LB media in deep-well plates and grown overnight. Cells are pelleted by centrifugation at 4000*g* for 30 minutes and suspended in P1, P2 and P3 buffers (from Zymo) in a 1 : 1 : 2 ratio. Plates are centrifuged at 4000*g* for 30 minutes and the supernatant is transferred carefully into 3 times the volume of absolute isopropanol to precipitate the plasmid. Plates are again centrifuged at 4^◦^C for 1 hour at 1800*g* and isopropanol is decanted and the plates air-dried at room temperature for 3 − 4 hours. 50*μ*L of nuclease-free is added to each well and the plates are incubated in 42^◦^C water for 10 minutes to dissolve the pelleted plasmid. The plates are then incubated on ice with the chemically competent cells of the corresponding TF Library strains for 30 minutes, heat-shocked, and recovered in SOC for 1 hour. Plates are centrifuged again and the pellet is resuspended in 30*μ*L SOC and patterned column-wise in LB-kanamycin plate using an 8-channel multi-channel pipette, plates are incubated overnight and individual colonies are obtained.

#### Growth and FACS measurement

The control and library strains of a given TF carrying independent promoter variants are grown overnight in 300*μ*L LB supplemented with antibiotics, kanamycin, and carbenicillin (for library strains, chloramphenicol is also added). The strains are diluted in a ratio of 1 : 5000 in 300*μ*L M9-minimal media supplemented with glucose in a 2 mL deep-well plates and grown at 37^◦^C to an OD of 0.2 − 0.5. The theoretical prediction in Fig. 1c is valid for saturated TF concentrations, effectively removing TF concentration as a variable. However, achieving true saturation may be challenging, particularly for weak binding sites (Fig. S1c). In this study, we approximate fully induced TF concentration (*T F*^++^) as equivalent to saturated levels. This assumption does not significantly impact our conclusions, provided that the TF concentration stays constant and relatively high (SI text 1.2 and Fig. S1d, high TF regimes). Except for SoxS, other TFs are induced at 25 ng/mL anhydrotetracycline (aTC). SoxS is induced with 6 ng/mL aTC as inducing SoxS to higher concentration reduced the growth of cells significantly. For MetR, the comparison is made between uninduced library strain and 25 ng/mL aTC as the MetR knockout cannot grow in M9-minimal media. For physiological perturbations, M9-minimal media is supplemented with different carbon sources (glycerol, galactose, L-arabinose, sodium pyruvate, or sodium acetate) to achieve different growth rates. Measurements for sigma factor variation are performed similarly to that for the synthetic promoters in M9-minimal media with glucose and 25 ng/mL aTC. Vanillic acid is serially diluted two-fold starting from 10*μ*M concentration to achieve different levels of σ^28^ concentration. The strains were diluted 1 : 80 in M9-minimal media with no carbon or nitrogen source to arrest cells in a steady state and incubated on ice until measurements. The plates are measured for mCherry and YFP fluorescence using BD LSRFortessa with an HTS (*X* −20−*Model* : 656385) and using the default settings for high-throughput sample readout. The mCherry is measured using a PE-CF594 laser at 600V. For the YFP signal, it is important to choose a FITC voltage to accommodate the range of expression that is expected of the different mutants. We measured auto-fluorescence and the weakest promoters at different voltages of FITC to find the minimal voltage that could be used to differentiate the auto-fluorescence signal from the weakest signal (S6a,b). We measured all YFP fluorescence with the FITC at 300V. The CST calibration is performed every day.

#### Data analysis

The raw data is extracted from the ‘fcs’ files with a custom-built Matlab code and unsupervised gates are applied as described in (*27*). Mean and standard error is calculated for both mCherry and YFP fluorescence after subtracting the signal from the autofluorescence sample. Any sample with less than 5000 events is excluded from the analysis (except for acetate the events less than 1000 events are excluded). Fold-change is calculated by taking the mean of the ratio of fluorescence from the library strain to the control strain in 2 independent experiments. The data is then fit to the model with *αβ* and *β* as fit parameters and C_max_ set by the promoter mutant with the maximum fluorescence for a given TF. Cells with negative fluorescence are excluded only for visual purposes when plotting the histograms of single-cell distribution of fluorescence.

#### Measuring translation differences for native promoters of SoxS

The highest constitutive mutant is selected from the promoter library for decRp, fldAp, and poxBp. The strains are grown as described above in multiple 300*μ*L wells, pooled at steady state (to a total of 1mL), OD quantified, and cell pellets were frozen overnight. Total RNA is isolated using the monarch RNA purification kit from NEB following the manufacturer’s protocol. The resulting RNA is quantified using nanodrop and 2 ng/*μ*L of total RNA is used in a single-step RT-qPCR reaction with NEB’s Luna RT-reagent. Separate reactions were set for the *yfp* and the *kanamycin resistance* gene. Both genes are expressed from the same reporter plasmid and the kanamycin resistance gene readout is used as a housekeeping gene control for normalization. Reactions were set up with two biological and two technical replicates each. Standard curves for both *kanamycin* gene and *yfp* are prepared using the 6 dilutions of the plasmid. The concentration of *yfp* mRNA level is first normalized with the concentration of *kanamycin* mRNA level and then by the total RNA yield per OD of the cell. This factor is then used to correct all the constitutive expressions of the corresponding promoters.

### Supplementary Text

#### Effect of changes to C_max_

The choice of *C*_max_ affects *α* and *β*, as the three variables are interdependent and difficult to decouple. Despite identical RBS and TSS, differences in the 5^′^-UTR between upstream and downstream promoters (due to the downstream TF binding site) result in translational variability for the same basal promoter (Fig. S1e). Therefore, we did not use a universal *C*_max_ for all 8 synthetic constructs. However, any choice of *C*_max_ only results in larger values of *β* (and does not yield smaller values), while leaving the product *αβ* unchanged (See SI Fig. S1f). In each case, the scaling relationship conforms to the prediction from the stabilizing model (*β* > 1) that fold-change scales as ∼ *C*^−1^ over most of the range. This data corresponds to the limit where (*β* − 1)*c*/*C*_max_ ≫ 1.

#### Thermodynamic model for constitutive expression versus finite TF numbers

The complete thermodynamic model of gene regulation is derived in (*14,15*) and shown in equation 4 below:

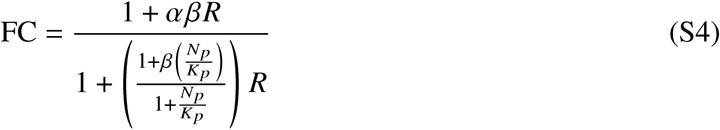

The promoter strength, 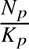 is related to the constitutive expression as shown by equation 1. Replacing 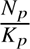 of equation 4 with the relationship shown in equation 1 simplifies as follows

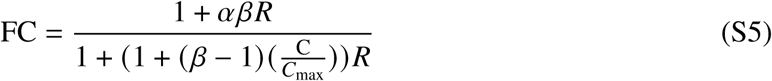

Fold-change predicted from the above expression for a certain value of *α* and *β* with varying concentrations of R is shown in Fig. S1d. For *β* > 1, as the TF number increases the curves spread out to the linear regimes for both activators (*α* = 2) and for repressors (*α* = 0.01). When R is infinite, 1 <<< *R*, the equation in 6 reduces to the equation in 3 (in the main text).

#### TF concentration during the measurements for physiological perturbations

Changes in carbon sources will affect global cellular physiology which means the levels of the TFs will also be impacted. As shown in SI text 1.2. and Fig. S1d, the effective concentration of TF has to be infinite for the fold-change to be independent of R. Effective concentration is a convolution of two factors: concentration of the TF and the affinity of the binding site for the TF. We know that the binding site for LacI used here (O1 binding site) has a very high affinity for LacI (*26*) whereas the binding site for CpxR is relatively weak (*10*). Thus, repression by LacI is very strong and attains maximum fold-change at mCherry values of 10^3^ fluorescence units Fig. S1c. However, regulation by CpxR is still not saturated in the range of aTC used in this study, and care is taken to achieve a uniform concentration of mCherry in different media Fig. S2c. Despite that, the mCherry levels (and hence the regulation) in pyruvate and acetate minimal media are higher than in other carbon sources for CpxR Fig. S2b. Hence, these two media are excluded from the figure in the main text. However, measurements in pyruvate and acetate for CpxR still show the same inverse scaling but at a higher fold-change value (shift in ‘y’ values) than that for other media (Fig. S2b). This is consistent with the behavior at varying finite TF concentrations illustrated in Fig. S1d.

#### Influence mCherry tag on the TF function

We wanted to exclude any possibility of tagging TFs with mCherry to interfere with its function. So, we wanted to test if the stabilizing behavior observed for the TF holds good irrespective of the mCherry tag. We chose LacI as it had the highest value for *β* and measured the regulation in strains expressing an untagged version of the TF. As shown in Fig. S2a, we did not see any difference in the regulation for strains expressing either the tagged or untagged version of our TF.

---

**Figure S1:**
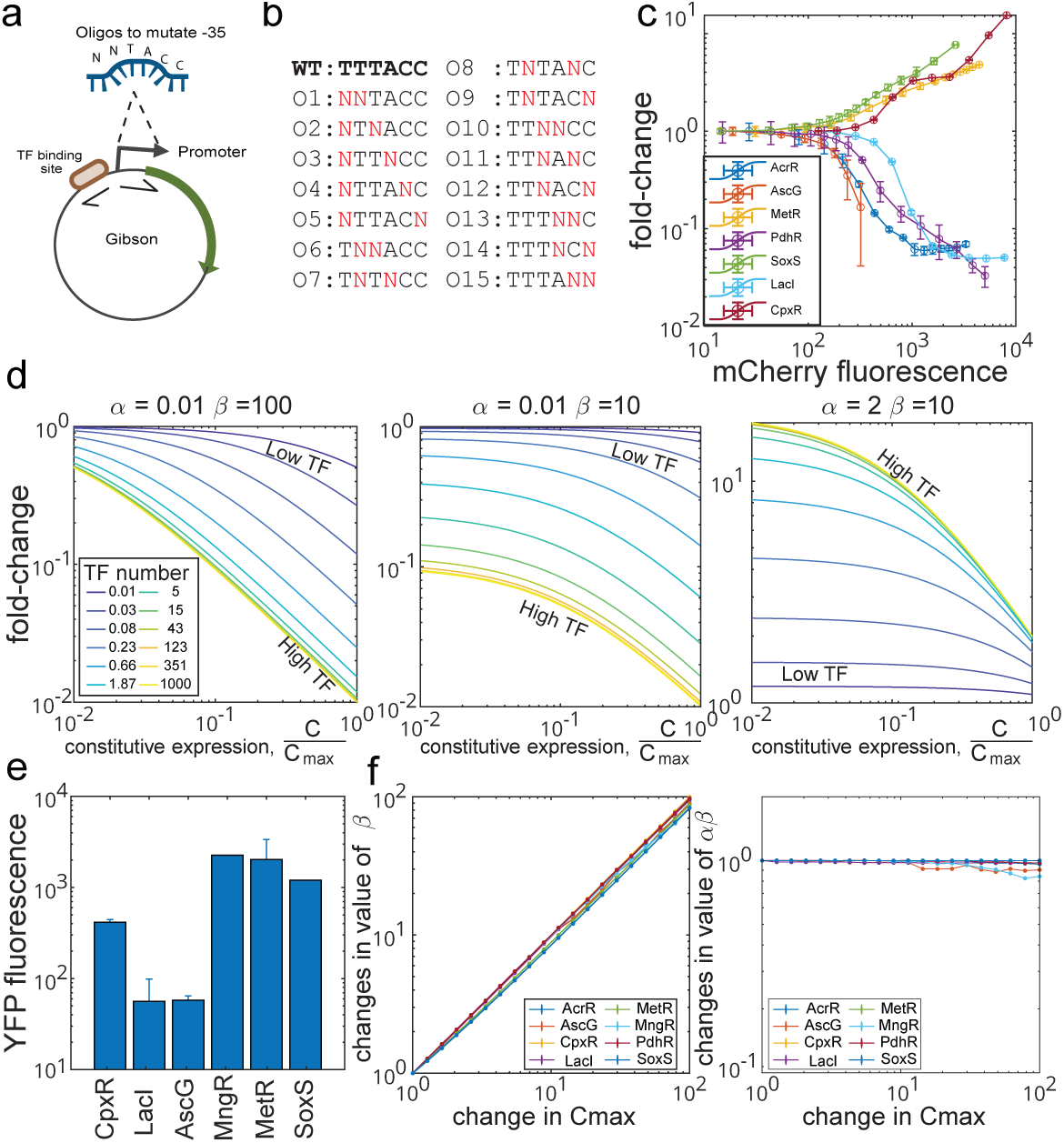
Synthetic promoter constructs: a) Introducing oligos with mutations to replace the −35 element of DL5p. b) Oligo pools to systematically change two nucleotides of the −35 sequence resulting in a combination of 153 unique −35 elements. c) Plot of mCherry fluorescence (achieved by titrating aTC) versus fold-change for the TFs shown in Fig. 2c. Plot indicates that two repressors (LacI, and AcrR) have achieved saturated fold-change whereas the three activators (CpxR, MetR and SoxS) and one repressor (PdhR) have still not achieved saturated fold-change for a range of aTC concentrations tested. d) Fold-change predicted for changing constitutive expression at finite TF concentration. e) Bars represent the YFP fluorescence value for the native DL5p with different TF binding sites. f) Plot showing the impact of changes to the *C*_max_ on the fit parameters *β* (left) and *αβ* (right)

**Figure S2:**
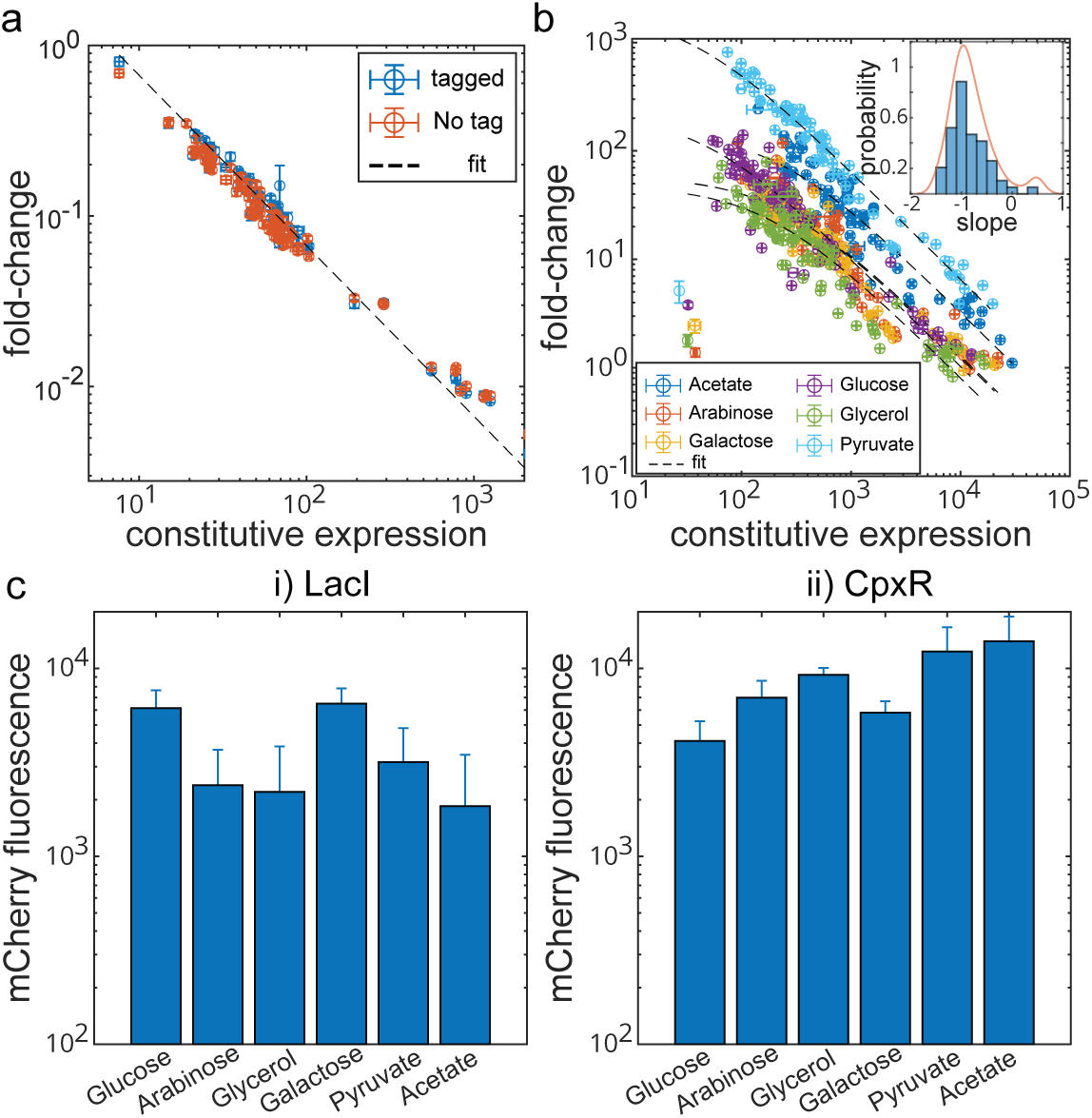
Altering cellular physiology. a) Comparing the impact of the mCherry fusion to the TF on the relationship between constitutive and regulated expression. Strains expressing the un-tagged version of LacI (red circles) is transformed with LacI promoter mutants and the regulated expression is measured as described above. b) Plot as in Fig. 3a excluding the data for LacI and including the data for CpxR measured in acetate and pyruvate. The insert shows the distribution of the slope of each promoter mutant across all 6 media for CpxR. c) Bar graph of mCherry expression in 6 different carbon sources for LacI (i) and CpxR (ii). mCherry levels for LacI are above the threshold where promoters are saturating however, CpxR is still not in the saturating regime and hence there is more spread in the data.

**Figure S3:**
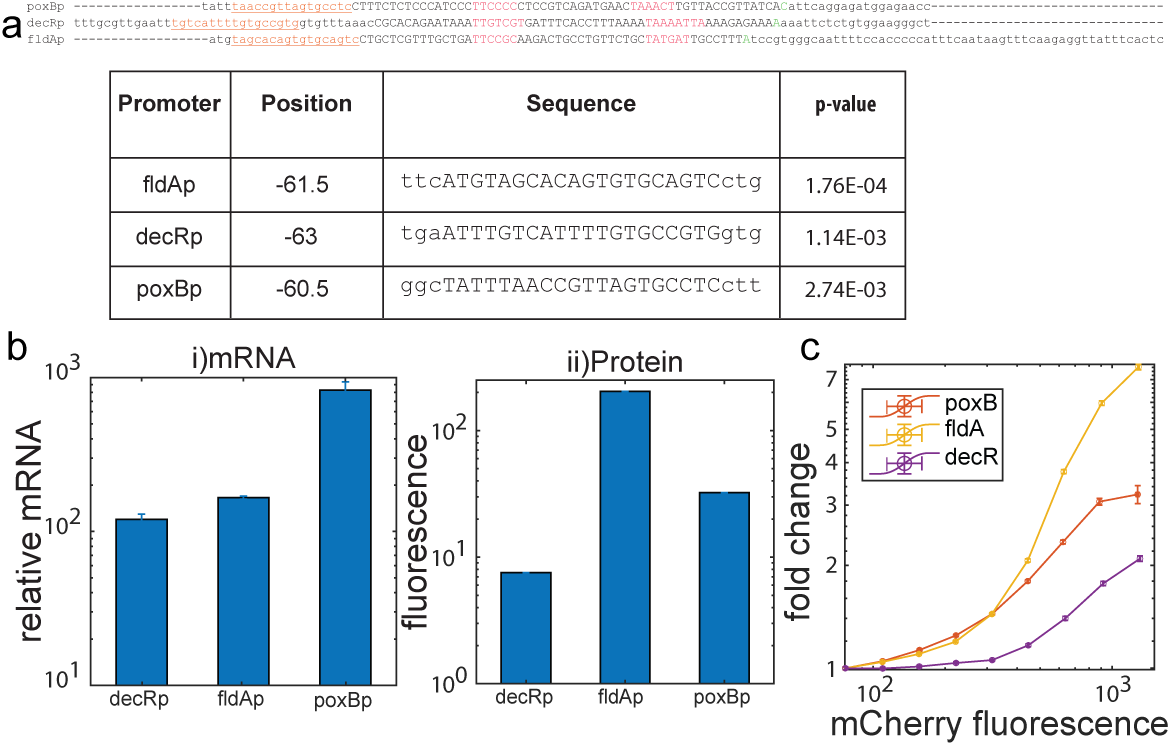
Regulation by SoxS. (a) Top panel shows the sequence of promoters used in Fig. 4a. The binding site is underlined and in brown. The −35 and −10 elements are in red. The TSS is in green. The bottom table lists the p-values of the binding sites in the TF PSSM browser of regulonDB. A list of all the sequences of SoxS used in this analysis is in the supplementary data file (b)Relative mRNA (i) and protein concentration (ii) of the three native promoters used to correct for the constitutive expression in Fig. 4a. (c) Regulation of the three native promoters of SoxS at different TF concentrations.

**Figure S4:**
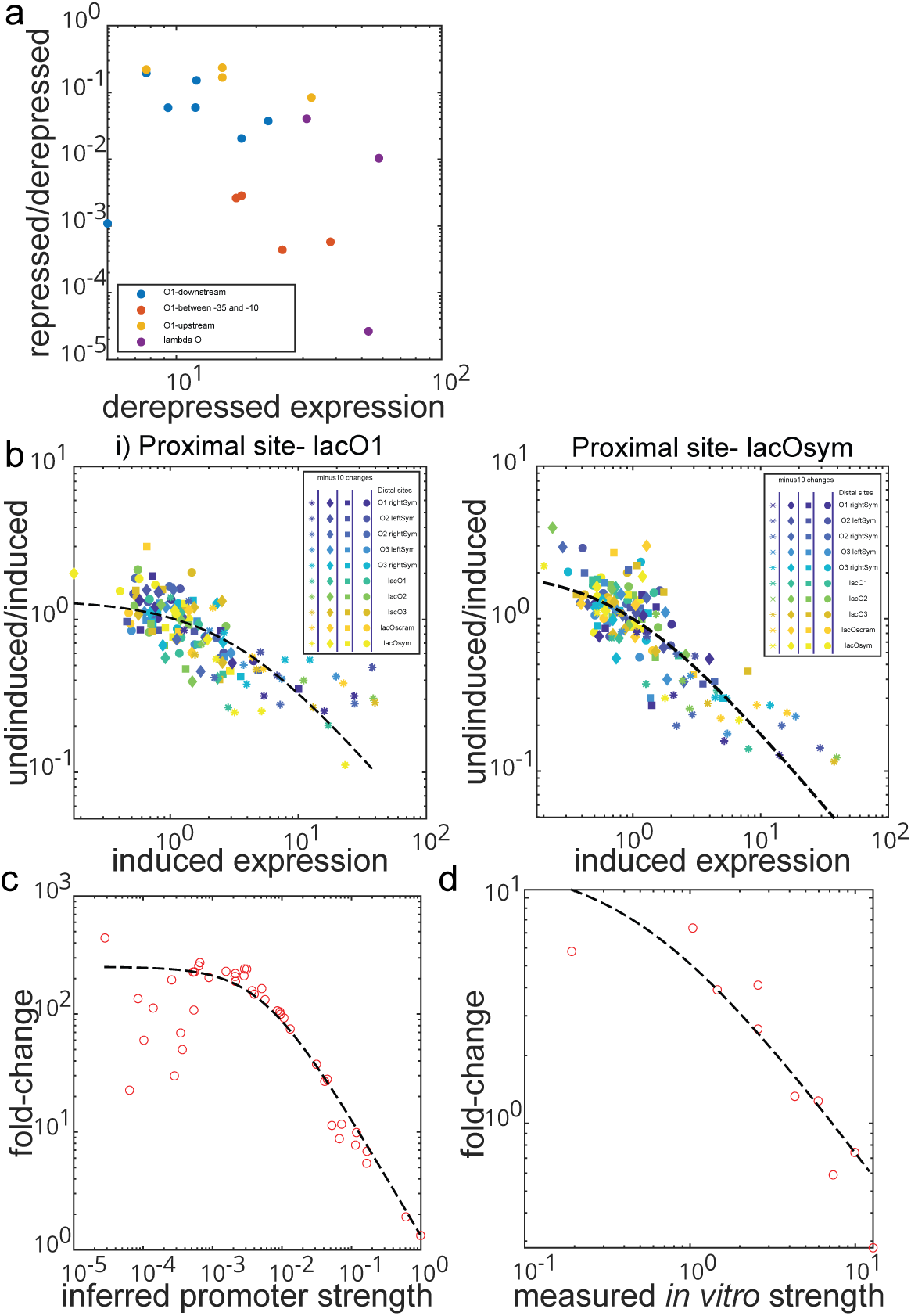
Raw data from external sources used in the study. (a) Data for LacI regulation measured by (*21*) (b) Data for LacI from (*43*). (c) Data for AraC from (*44*) (d) Data for CarD from (*46*)

**Figure S5:**
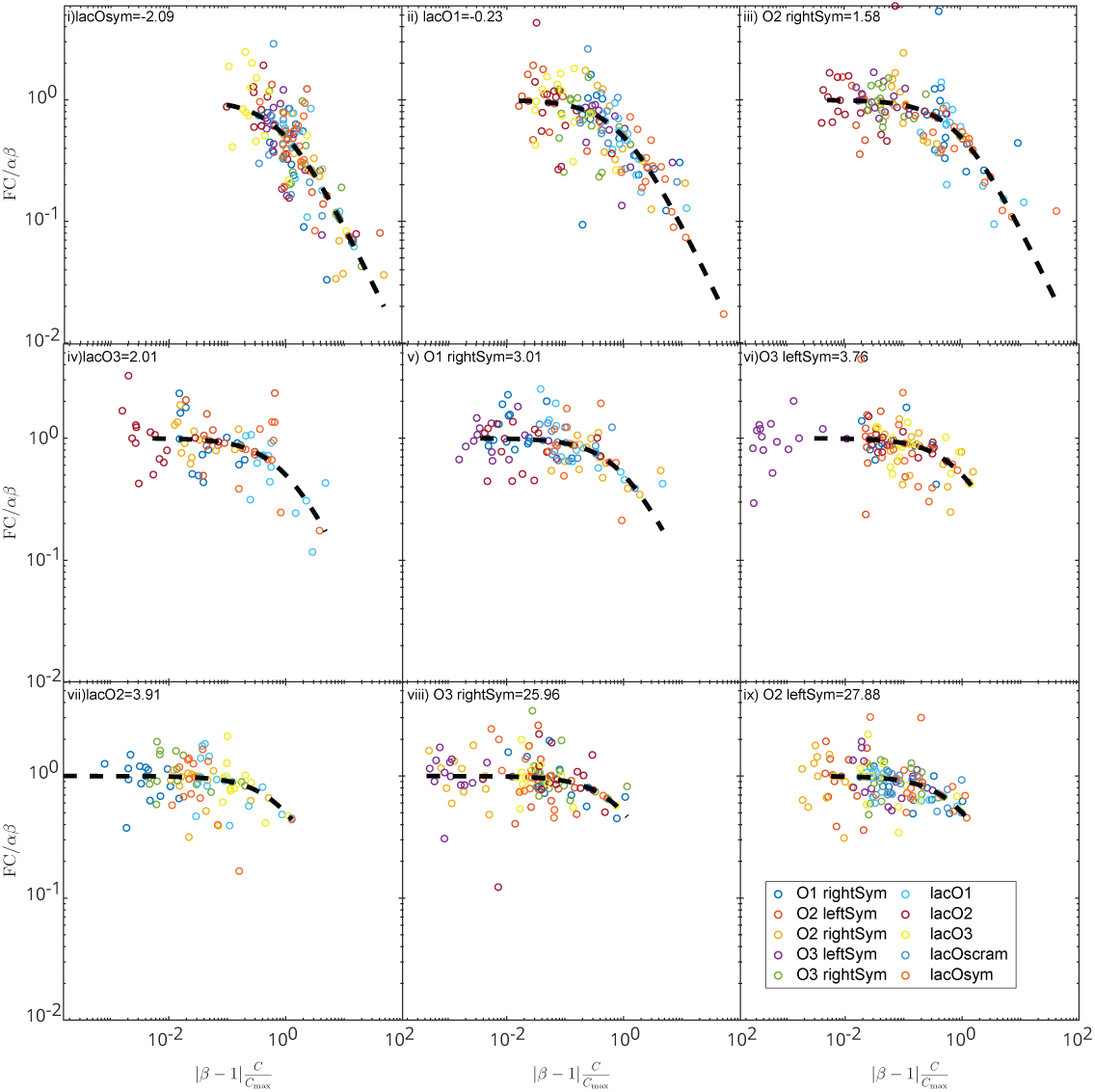
Looping data for all sites from Yu *et al* (*43*). Data in each plot have the same proximal site and 10 different distal sites. The plots are arranged in increasing order of the inferred affinity of the distal site. The weaker the distal site is the weaker the regulation.The constitutive expression in this data set is achieved by using IPTG to inactivate LacI whereas in our dataset it is the complete knockout of the TF.

**Figure S6:**
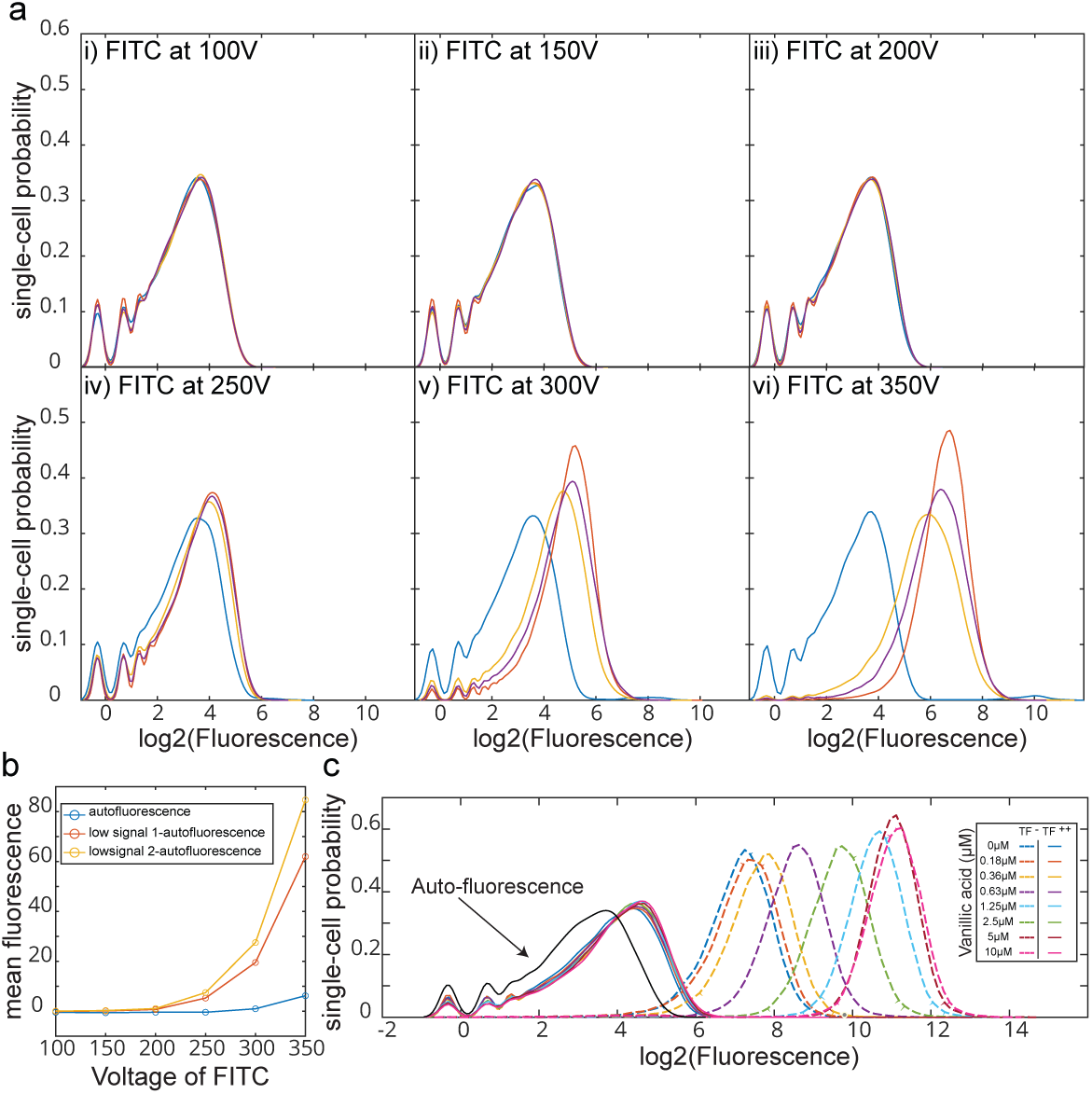
Choice of FITC voltage. (a) Probability density function of single-cell fluorescence of auto-fluorescent strain and strains expression lowest yfp expression measured at different voltages of FITC. (b) Blue data points correspond to the mean of fluorescence across different voltages for auto-fluorescence strain. The yellow and red data points are the mean of the fluorescence upon auto-fluorescence subtraction. All data points in this paper are measured at 350 Voltage in the FITS channel to be best able to differentiate lower signals from auto-fluorescence and still have the highest signals from the non-saturating regime. (c) plot as in 5c with the auto-fluorescence data included.

**Figure S7:**
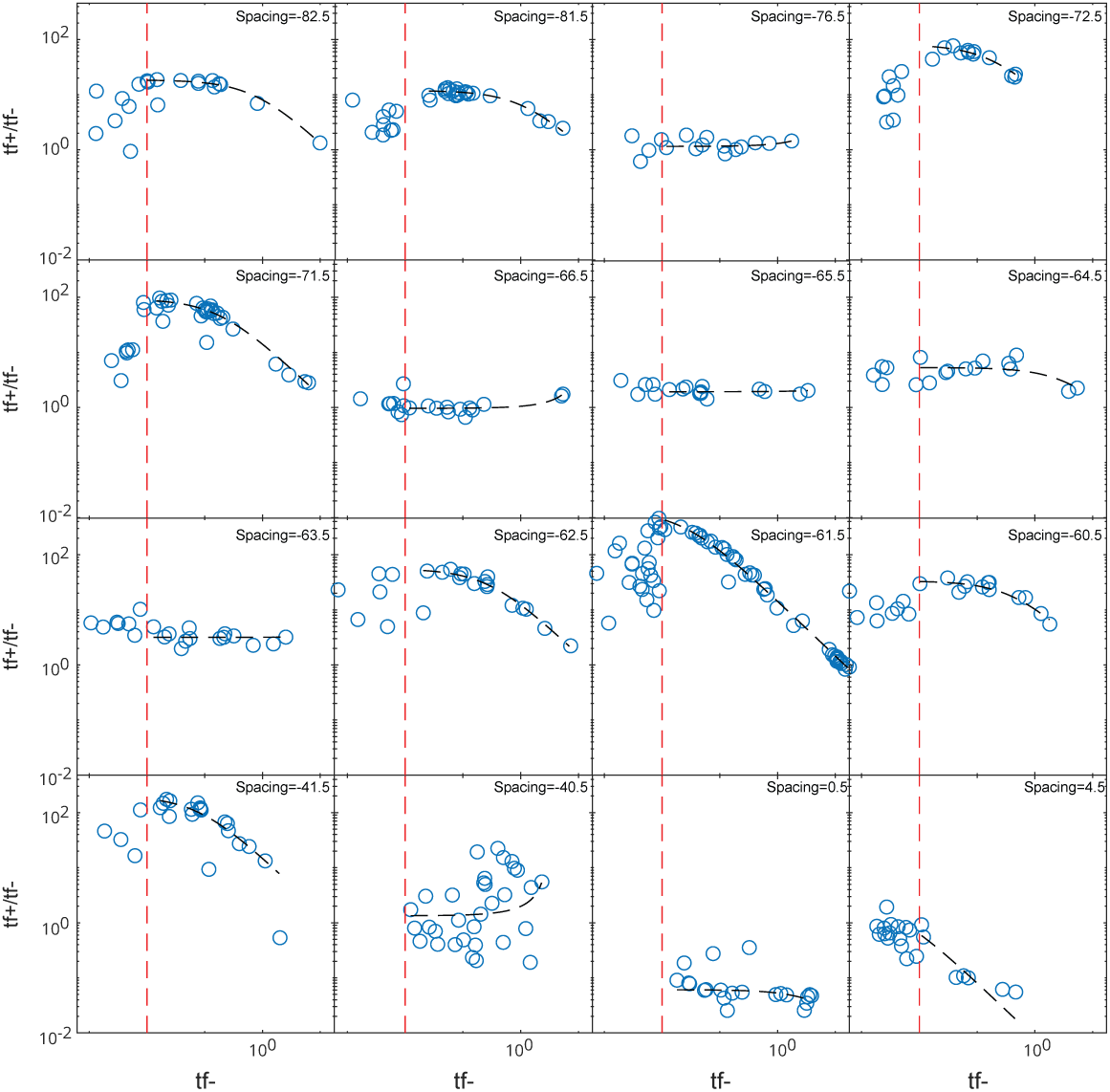
Regulation by Crp data by Forcier et. al. (*45*). Plots showing the ratio, 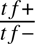 (which is analogous to fold-change) to *t f* − (analogous to constitutive expression in our data) for Crp at different spacing measured by LacZ activity assay. Spacing is the distance between the TSS on the promoter to the middle of the binding site for Crp. A noteworthy point is that *t f* − in Forcier et. al. (*45*) is a measurement without cAMP with the assumption that Crp is inactive without its co-factor, cAMP. The data that falls below the red dashed lines in each plot is excluded from our analysis to account for background and sensitivity in their data.

**Table S1:**
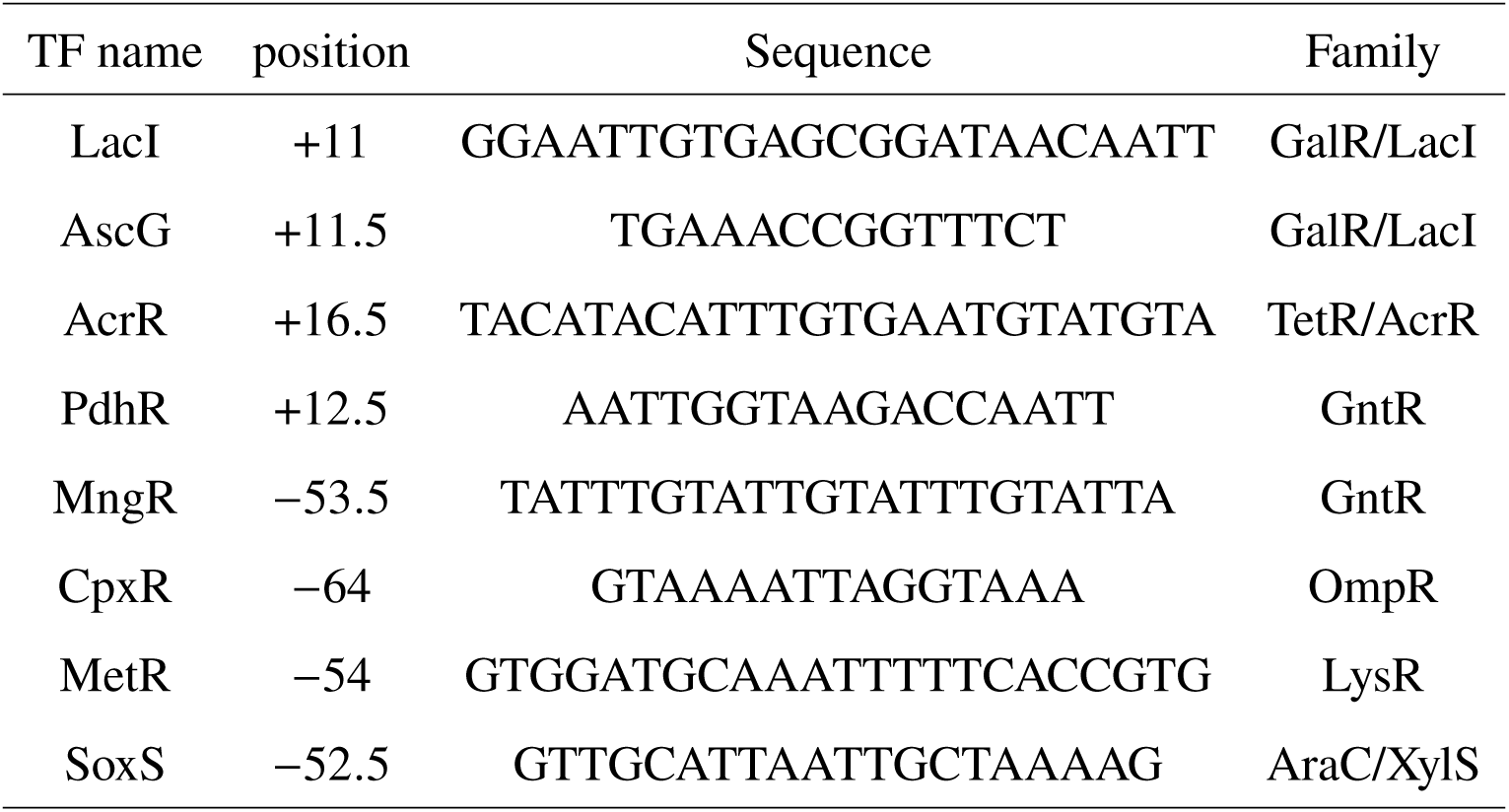
Sequence list for TF binding sites. Information of TF binding site identity, position and sequence for each TF binding site from Fig. 2.

